# The triple combination of Remdesivir (GS-441524), Molnupiravir and Ribavirin is highly efficient in inhibiting coronavirus replication in human nasal airway epithelial cell cultures and in a hamster infection model

**DOI:** 10.1101/2024.05.14.594200

**Authors:** Thuc Nguyen Dan Do, Rana Abdelnabi, Bernadett Boda, Samuel Constant, Johan Neyts, Dirk Jochmans

## Abstract

The use of fixed dose-combinations of antivirals with different mechanisms of action has proven a key in the successful treatment of infections with HIV and HCV. For the treatment of infections with SARS-CoV-2 and possible future epi-/pandemic coronaviruses, it will be important to explore the efficacy of combinations of different drugs, in particular to avoid resistance development, such as in patients with immunodeficiencies. As a first effort, we studied the antiviral potency of combinations of antivirals. To that end, we made use of primary human airway epithelial cell (HAEC) cultures grown at the air-liquid interface that were infected with the beta coronavirus OC43. We found that the triple combination of GS-441524 (parent nucleoside of remdesivir), molnupiravir, and ribavirin resulted in a more pronounced antiviral efficacy than what could be expected from a purely additive antiviral effect. The potency of this triple combination was next tested in SARS-CoV-2 infected hamsters. To that end, for each of the drugs, intentionally suboptimal or even ineffective doses were selected. Yet, in the lungs of all hamsters that received triple prophylactic therapy with suboptimal/inactive doses of GS-441524, molnupiravir, and ribavirin, no infectious virus was detectable. Our finding indicate that co-administration of approved drugs for the treatment of coronavirus infections should be further explored but also against other families of viruses with epidemic and pandemic potential for which no effective antiviral treatment is available.

## INTRODUCTION

Three novel (beta)coronaviruses emerged during the past two decades that are highly pathogenic to man. These are the severe acute respiratory syndrome coronavirus (SARS-CoV) in 2002, the Middle East respiratory syndrome coronavirus (MERS-CoV) in 2012 and SARS-CoV-2 in 2019. The COVID-19 pandemic, that was caused by SARS-CoV-2, resulted in ∼7 million confirmed and ∼28 million estimated deaths so far (https://ourworldindata.org/explorers/coronavirus-data-explorer). During and after the emergence of SARS-CoV in 2002, there have been efforts to develop inhibitors of the viral main protease (Mpro or 3CLpro). Twenty years later, this work laid the basis for the rapid development of nirmatrelvir, a potent Mpro inhibitor with pan-coronavirus coverage (1). Remdesivir and molnupiravir are both prodrugs of nucleoside analogues that target, as their 5’-triphosphate metabolite, the viral RNA-dependent RNA polymerase (RdRp) (2) or induce error catastrophe (3), respectively. These molecules were originally developed for the treatment of infections with other viruses but are also endowed with broader spectrum antiviral activity, including activity against coronaviruses. These drugs were therefore repurposed for the treatment of infections with SARS-CoV-2 (2), (4). Remdesivir requires intravenous administration, but an oral form, obeldesivir (5), is currently in clinical development for the treatment of COVID-19. GS-441524 is the parent nucleoside of both remdesivir and obeldesivir. Intracellularly, these molecules (GS-441524, remdesivir and obeldesivir) are converted to the tri-phosphate form of GS-441524, which is incorporated in the viral RNA (vRNA) and acts as a chain terminator (5). Molnupiravir, is an oral broad-spectrum antiviral designed to combat alphavirus and influenza viruses (6) and is in some countries conditionally approved for the treatment of COVID-19 (7). Molnupiravir is intracellularly converted to its 5’-triphosphate form and incorporated in the vRNA leading to lethal mutagenesis and hence inhibition of viral replication (7). Ribavirin is a guanosine analogue with broad-spectrum antiviral activity that has been widely used to treat hepatitis C virus (HCV) infection, to some extent and with limited success for the treatment of RSV infections, and infections with viral haemorrhagic fever viruses. Although ribavirin has been explored for the treatment of COVID-19, no significant efficacy has been shown (8), (9). Several mechanisms of action for ribavirin have been proposed such as depletion of intracellular nucleoside pools, inhibition of viral RNA capping, direct inhibition of the viral RNA polymerase, inducing of viral mutagenesis, and modulation of host immune responses (for review see (10)).

The high conservation of viral RdRp, the lack of host homologs, and the relative high barrier to resistance make broad spectrum RdRp inhibitors ideal candidates for antiviral therapy. Additionally, combination treatment with antivirals that possess different mechanisms of action proves to be superior to monotherapy in controlling viral infections caused by the human immunodeficiency virus (HIV) and the hepatitis C virus (HCV). Such fixed-dose combinations also suppress or delay the onset of drug-resistance development when compared to single agents (11). The use of combinations of antiviral drugs against SARS-CoV-2, may also result in a more potent effect than of monotherapy. In addition, it may allow for dose-reduction and a reduced risk of resistance development. We previously reported that favipiravir significantly potentiates the antiviral efficacy of molnupiravir when co-administered to SARS-CoV-2 infected Syrian hamsters (12). Similarly, the combined treatment of SARS-CoV-2-infected Syrian hamsters with favipiravir and GS-441524 has been reported to be more efficient compared to the monotherapies (13). Recently, it has also been reported that the combination of suboptimal concentrations of molnupiravir and remdesivir completely blocks the production of SARS-CoV-2 infectious virus particles in the apical wash of human nasal airway epithelium cultures (14). We also demonstrated that the combination of molnupiravir and GS-441524 is highly effective in reducing SARS-CoV-2 replication/infectivity in a hamster infection model (15). We here explore a next step in these combination studies by studying the efficacy of the triple combination of GS-441524, molnupiravir, and ribavirin against coronavirus infection in primary human airway epithelial cells (HAEC) infected with the beta coronavirus OC43 and in Syrian hamsters infected with SARS-CoV-2.

## MATERIALS AND METHODS

### Virus isolates

Human coronavirus OC43 (HCoV-OC43) was isolated directly on HAE from nasal pharyngeal swabs as previously described (16). The HCoV-OC43 virus stock was then produced on HAE culture by collecting and pooling apical washes (17).

The SARS-CoV-2 variant of concern, B.1.351 (EPI_ISL_896474|2021-01-11 or beta variant) was named according to its own lineage. The strain was isolated from a nasopharyngeal swab of a RT-qPCR confirmed patient. The generation of virus stock was fully reported elsewhere (18).

All SARS-CoV-2 infectious virus-related work was carried out in a biosafety level 3 (BSL-3) facility at the Rega Institute for Medical Research, KU Leuven, according to institutional guidelines.

### Compounds

GS-441524 and ribavirin were purchased from Carbosynth (United Kingdom) while molnupiravir (EIDD-2801) was purchased from Excenen Pharmatech Co. Ltd. (China). In *ex-vivo* experiments, 10 mM stock solution of each compound was prepared in analytical grade dimethyl sulfoxide (DMSO, Sigma). For *in vivo* study, GS-441524 was formulated as a 15 mg/mL stock in a vehicle containing 30% PEG-400 (Sigma) and 1% DMSO in PBS. Molnupiravir was formulated as 50 mg/mL stock in 10% PEG-400 and 2.5% Kolliphor-EL (Sigma) in water. Ribavirin was formulated as 15 mg/ml solution in PBS.

### Viral infection of human airway epithelial cells

Human nasal airway epithelial cells (HNAEC or MucilAir, cat. no. EP01MD) from a pool of fourteen different donors were obtained from Epithelix (Switzerland) in an air-liquid interphase (ALI) setting and cultured at 37°C under standard conditions (5% CO2, 100% humidity). Through the course of the experiment itself the cultures were maintained at 34°C. Prior to infection with HCoV-OC43 (3×10^5^ viral copies per insert), the apical site of the cultures were washed once with pre-warmed MucilAir medium (Epithelix, cat. no. EP04MM) and the inserts were then transferred to medium with or without compound. After 1 hour pre-incubation, 100 µL of HCoV-OC43 was added at the apical site, incubated for 1.5 hours, and the unbound virus was then removed by washing the apical site three times with pre-warmed basal medium. The last of these apical washes was used for quantification of virus background on day 0. Also on day 1, 2, 3, 6, 8, and 10, an apical wash was collected for virus quantification. On day 2 and 4, the basolateral medium of the HAE culture was refreshed with medium containing compound (or without compound for the controls). On day 6 and 8, the basolateral medium was refreshed with medium without compound for all conditions. Collecting washes was continued for four days after the treatment regimen. Samples were stored at −80°C until analysis.

### SARS-CoV-2-hamster infection model

The hamster infection model of SARS-CoV-2 has been described before (18). Female Syrian hamsters (*Mesocricetus auratus*, Janvier Laboratories) were housed as pairs in individually ventilated isolator cages (IsoCage N Bio-containment System, Tecniplast) at 21°C, 55% humidity, and 12:12 day/night cycles. Housing conditions and experimental procedures were approved by the ethics committee of animal experimentation of KU Leuven (license P065-2020). For infection, female hamsters of 6-8 weeks old were first anesthetized with ketamine/xylazine/atropine and inoculated intranasally with 50 µL containing 1×10^4^ TCID_50_ SARS-CoV-2 beta variant (day 0). After four days of infection, animals were euthanized for collection of the lungs and further analysis by intraperitoneal (i.p.) injection of 500 μL Dolethal (200 mg/mL sodium pentobarbital).

The doses of compounds were selected such that they result, as monotherapy, in a limited antiviral activity (i.e. < 1 log_10_ reduction in infectious virus titers). The selection of the molnupiravir dose (75 mg/kg, BID) was based on a dose-response study against the wild-type strain that we published previously (19). For GS-441524, we previously tested both 25 and 50 mg/kg BID in our model. The 25 mg/kg BID dose did not result in any reduction in infectious viral loads (unpublished data) while the 50 mg/kg BID dose reduced viral RNA and infectious virus titers in the lung by 1.2 log_10_ copies/mg and 0.5 log_10_ TCID_50_/mg lung tissue, respectively (15). Therefore we selected GS-441524 at 25 mg/kg BID for the current study. For ribavirin, we explored 25 and 50 mg/kg BID in a pilot study. Both resulted in no antiviral efficacy. However, the 50 mg/kg BID dose resulted in weight loss in the treated hamsters, therefore we selected the 25 mg/kg BID for the current study. Hamsters were treated BID via oral gavage with either vehicle (n=12), molnupiravir 75 mg/kg (n=10), GS-441524 25 mg/kg (n=10), ribavirin 25 mg/kg (n=10) or combination of the three compounds at the same doses as selected for monotherapy (n=12) starting from the time of infection (d0) with SARS-CoV-2 beta variant. Smaller groups of animals were also treated with the double combination of either molnupiravir+GS-441524 (n=6), molnupiravir+ribavirin (n=4) or GS-441524+ribavirin (n=4). The BID treatments were done 8 h apart and the compounds were given sequentially for the combination therapy. All the treatments continued until day 3 post-infection (p.i.). Hamsters were monitored for appearance, behavior and weight. At day 4 p.i., hamsters were euthanized as mentioned earlier. Lungs were collected for viral loads quantification. Two independent experiments were done.

### End-point titration assay

Hamster’s lung tissues were homogenized using bead disruption (Precellys) in minimal essential medium (MEM) and centrifuged (10000 rpm, 5 min, 4°C) to pellet the cell debris. Serial dilutions of homogenized supernatant were performed on confluent Vero-E6 cells in 96-well plates to quantify infectious SARS-CoV-2 particles based on the absence or presence of virus-induced cytopathic effect (CPE). Viral titers were calculated by the Reed and Muench method (20) and expressed as TCID50 per milligram of tissue.

### RNA extraction and quantitative reverse transcription-PCR (RT-qPCR)

Viral RNA of HCoV-OC43 from the apical washes were extracted using QIAamp viral RNA kit (Qiagen, cat. no. 52906) according to the manufacturer’s instructions. In parallel, a ten-fold serial dilution of corresponding virus stock was extracted. The amount of viral RNA expressed as number of copies per millilitre (copies/mL) for HCoV-OC43. Viral RNA was quantified by RT-qPCR using QuantiTect Probe RT-PCR (Qiagen, cat. no. 204445) as previously reported (17).

Viral RNA of SARS-CoV-2 was extracted from homogenized hamster lung tissues using bead disruption (Precellys) in 350 µL TRK lysis buffer (E.Z.N.A. total RNA Kit, Omega Bio-tek) followed by a centrifugation (10000 rpm, 5 min) according to the manufacturer’s protocols. RT-qPCR was executed on a LightCycler thermocycler (Roche) using iTaq universal Probes One-step RT-qPCR kit (BioRad) with N2 primers and probes (IDT) as previously described (21). A serial dilution of SARS-CoV-2 cDNA standards (IDT) was used to express viral genome copies per mg tissue.

### Trans-epithelial electrical resistance (TEER)

The electrical resistance of a cellular monolayer is an indicator of the barrier integrity, measured by an epithelial voltohmmeter (EVOM3, World Precision Instruments). During the washing step, 250 µL fresh medium was added to the apical side of the HAE culture, incubated for 15 minutes at 34◦C, followed by the subsequent resistance measurement according to the manufacturer’s instructions. Briefly, the two electrodes of the “chopstick” were allocated in the apical and basal compartments. The resistance of the culture tissue was then subtracted (100 Ω) from the recorded value. The subtracted value (100 Ω) is the resistance of the semipermeable membrane without the cell layer, provided by manufacturer. TEER values were presented in units of ꭥ.cm^2^ and calculated as:

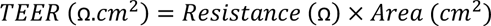

where Resistance and Area are the cell specific resistance and the surface of semipermeable membrane, respectively. The TEER within one specific day was normalized to the uninfected-untreated controls.

### Cytotoxicity measurement

Compound/Virus-induced toxicity was measured by a colorimetric assay to detect lactate dehydrogenase (LDH) released into the basolateral medium using CyQUANT LDH Cytotoxicity Assay kit (Invitrogen, cat. no. C20300). In short, 50 µL of each sample basal medium was transferred into a 96-well plateand mixed with 50 µL of reaction mixture. Samples were incubated at room temperature for 30 minutes, protected from light, followed by the addition of 50 µL of stop solution to each well. Medium from HAE cultures that were lysed by Triton X-100 10% overnight was used as maximal LDH activity control. In parallel, the fresh MucilAir medium served as spontaneous LDH activity control and LDH positive control were included in each experiment. The absorbance at 490 nm and 680 nm were measured. The 680-nm absorbance values were subtracted from 490-nm absorbance values to calculate the LDH activity. Percentage (%) cytotoxicity was determined according to following formula:

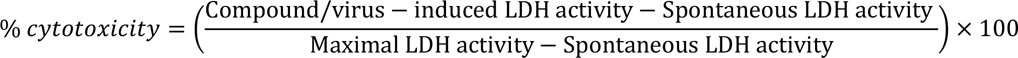

### Pharmacodynamic interaction analysis

The interactions within double and triple therapies (ribavirin + GS-441524, molnupiravir + GS-441524, ribavirin + molnupiravir, and ribavirin + GS-441524 + molnupiravir) were analysed with the Bliss independence zero-interaction theory. According to this pharmacodynamic theory, if three molecules (A, B, and C) do not interact, i.e. they act independently, then the effect E of their combined actions can be predicted using the probability law of independent events (22). This predicted percentage inhibition *E*_ABC,Pred._ is determined by the formula:

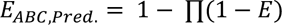

where ∏(1 − *E*) is the product of observed viral propagations of each single agent alone (i.e. *E_ABC,Pred._* = 1 − (1 − *E_A_*) × (1 − *E_B_*) × (1 − *E_C_*) = *E_A_* + *E_B_* + *E_C_* − *E_A_* × *E_B_* – *E_A_* × *E_C_* – *E_B_* × *E_C_* + *E_A_* × *E_B_* × *E_C_*). Of note, the percentage viral inhibition/reduction in the latter equation can be converted from log_10_ viral reduction: *P* = (1 − 10^-L^) × 100 where P is the percentage inhibition/reduction and L is the log_10_ viral reduction.

The observed combined percentage inhibition *E_ABC,Obs._* is then compared with *E_ABC,Pred._*. If *E_ABC,Obs._* > *E_ABC,Pred._*, the compounds act in a synergistic manner; if *E_ABC,Obs._* < *E_ABC,Pred._*, there is antagonism whereas *E_ABC,Obs._* = *E_ABC,Pred._*, indicates additivity.

### Statistical analysis

All statistical comparisons in the study were performed in GraphPad Prism 9 (GraphPad Software, Inc.). Statistical significance was determined using the ordinary one-way ANOVA with Dunnett’s multiple comparison test (*ex vivo* data) or the non-parametric Mann-Whitney U-test (*in vivo* data). P-values of <0.05 were considered statistically significant. In the figures “ns” indicates a p > 0.05. Asterisks indicate a statistical significance level of *p < 0.05, **p < 0.01, ***p < 0.001, ****p < 0.0001.

## RESULTS

### Antiviral effect of triple combination of GS-441524, Molnupiravir and Ribavirin in HNAEC cultures infected with HCoV-OC43

To explore the antiviral efficacy of combinations of two or three drugs [GS-441524 (the parent nucleoside of remdesivir), molnupiravir and ribavirin] human nasal human airway epithelial cell (HAEC) cultures were infected with HCoV-OC43. The kinetics of virus replication was monitored over several days through quantification of virus released at the apical site of the cultures. We first defined the effective and suboptimal concentration of GS-441524 and molnupiravir for inhibition of HCoV-OC43 replication in nasal HAEC cultures. Treatment with 10 µM GS-441524 results in a “cure” of the cultures since no viral rebound was observed when the therapy is stopped. At 1 µM of GS-441524, however, the viral load is only marginally reduced (**Figure 1A**). At 3 µM, molnupiravir treatment results in a significant inhibition of viral replication (up to 1.5 log_10_ vRNA reduction) whereas 1 µM is clearly suboptimal (**Figure 1B**). Based on these results, we selected for both GS-441524 and molnupiravir a concentration of 1 µM as suboptimal for a subsequent combination experiments in nasal HAEC cultures. Ribavirin, at 10 µM, has no measurable effect on HCoV-OC43 replication in HAEC cultures (**Figure 2**).

**Figure 1.**
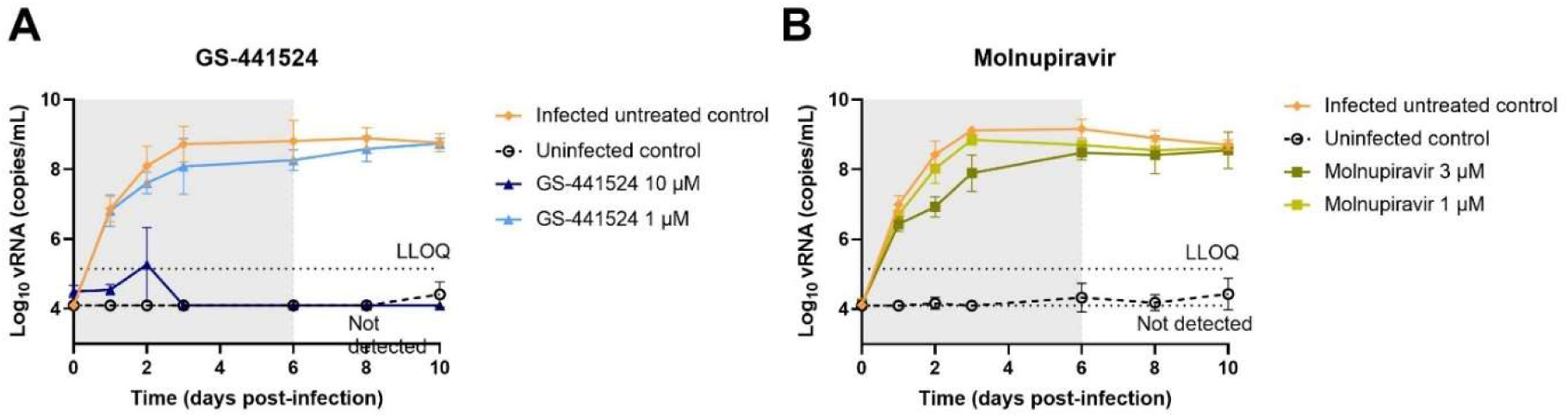
Antiviral activity of GS-441524 and molnupiravir against HCoV-OC43 in HAEC cultures. Compounds were added to the basal medium at different concentrations 1 hour prior to infection with HCoV-OC43 at 34°C. Basal medium, with or without compounds, was refreshed every other day from day 0 to day 6. Viral RNA in apical washes were quantified by RT-qPCR. Dose-response and time-dependent activity of GS-441524 (A) and molnupiravir (B). All data are mean ± SD of at least three replicates. Grey box indicates time of treatment. LLOQ presents lower limit of quantification.

**Figure 2.**
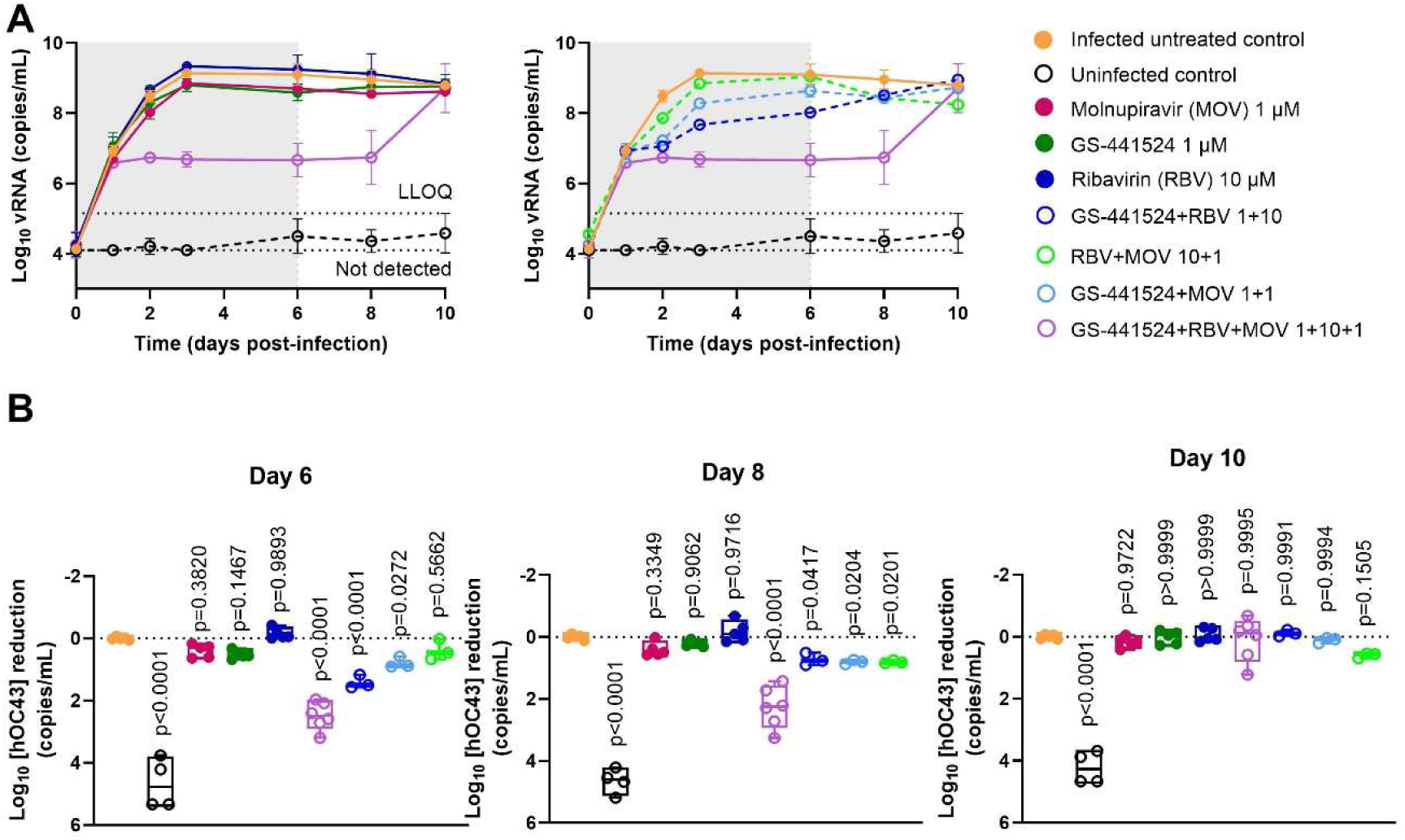
Comparing the antiviral activity of combinations of GS-441524, molnupiravir (MOV), and ribavirin (RBV) on HCoV-OC43 replication in nasal human airway epithelial cell (HAEC) cultures. Compounds were added to the basal medium starting 1 hour before infection and treatment continued for 6 days. Nasal HAEC were infected with HCoV-OC43 at 3×10^5^ copies/insert and incubated at 34°C. Viral RNA in apical washes was quantified by RT-qPCR. (A) Kinetics of HCoV-OC43 replication with or without different monotherapies and combinations of inhibitors. (B) Summaries of the antiviral activities of each treatment on day 6, 8, and 10 were analysed. Two independent experiments with nasal HAEC from a pool of donor were performed (n=3-6). LLOQ represents the lower limit of quantification. Statistical significance between infected untreated control and other groups was calculated by one-way ANOVA with two-sided Dunn’s post hoc test. Data are mean ± SD of at least three biological replicates.

Next, the effect of 1 µM GS-441524, 1 µM molnupiravir and 10 µM ribavirin, either alone or in double or triple combinations on HCoV-OC43 replication was assessed (**Figure 2A)**. Statistical analysis was performed for the results on day 6 p.i., which corresponds with the last day of treatment (**Figure 2B**). Neither the monotherapies nor the double combinations of either GS-441524 plus molnupiravir or molnupiravir plus ribavirin has a significant effect on virus replication. The combinations GS-441524 plus ribavirin results in a 1.4 log_10_ reduction in viral RNA but when combined with molnupiravir (thus GS-441524 plus ribavirin plus molnupiravir), 2.5 log_10_ RNA reduction in viral RNA is achieved. Upon cessation of therapy, virus replication increases again for both conditions.

The monotherapies result in less than 0.7 log_10_ (80%) vRNA reduction. As a conservative estimate we assume that each of the monotherapies reach 80% inhibition; using this estimate we compare the predicted effect of the triple combination with the observed effect. According to the Bliss model (described in the methods), the predicted inhibition would be 80% + 80% + 80% - 80%*80% - 80%*80% - 80%*80% + 80%*80%*80% = 99.2% (2.1 log_10_ vRNA reduction). The observed inhibition has a median of 99.7% (2.5 log_10_ vRNA reduction) with an interquartile range of 99.3% – 99.8%. These numbers, together with the fact that we started from an overestimation of the activities of the monotherapies (0.7 log_10_ vRNA reduction), indicate a synergistic interaction between GS-441524, molnupiravir, and ribavirin. Of note, there was no indication of toxicity in any of the conditions tested in trans-epithelial electrical resistance (TEER) measurement (i.e. determining the integrity and permeability of lung tissue) or when quantifying levels of released lactate dehydrogenase enzyme (**Supplementary data Figure S1**).

### The combined treatment of suboptimal doses of GS-441524 and molnupiravir with ribavirin efficiently inhibits SARS-CoV-2 replication in hamsters

We next studied whether the triple combination of GS-441524, molnupiravir and ribavirin results also in SARS-CoV-2 infected hamsters in a more than additive activity. Female hamsters were intranasally infected intranasally with 1×10^4^ TCID_50_ of SARS-CoV-2 (beta variant, lineage B.1.351) and were orally treated twice a day (BID) for four consecutive days starting one hour before infection. Doses of each of the drugs were in preliminary studies (see methods) determined to be suboptimal to inactive. In the current experiment, GS-441524 alone at 25 mg/kg/dose resultes in 0.2 log_10_ reduction of infectious virus titers in the lungs (at day 4 post infection), molnupiravir at 75 mg/kg/dose in 0.7 log_10_ reduction and ribavirin at 25 mg/kg/dose in no observable reduction of viral titers. None of the changes in infectious virus titers by the monotherapies was statistically significant. On day 4 p.i., infected hamsters from all groups exhibited < 10% weight change indicating that all treatments were well tolerated (**Figure 3**). The combination of GS-441524 with molnupiravir reduced infectious titers in the lungs by 0.6 log_10_ (p=0.72), compared to the vehicle control. No significant reduction in infectious titers in the lungs was observed in the ribavirin plus molnupiravir group. The combination GS-441524 plus ribavirin resulted in 0.80 log_10_ (p=0.51) reduction in infectious titers. Finally, in all 12 animals that had been treated with GS-441524 plus molnupiravir plus ribavirin, infectious titers in the lungs were reduced by > 3.7 log_10_ (p<0.0001) to undetectable levels (**Figure 3**).

**Figure 3.**
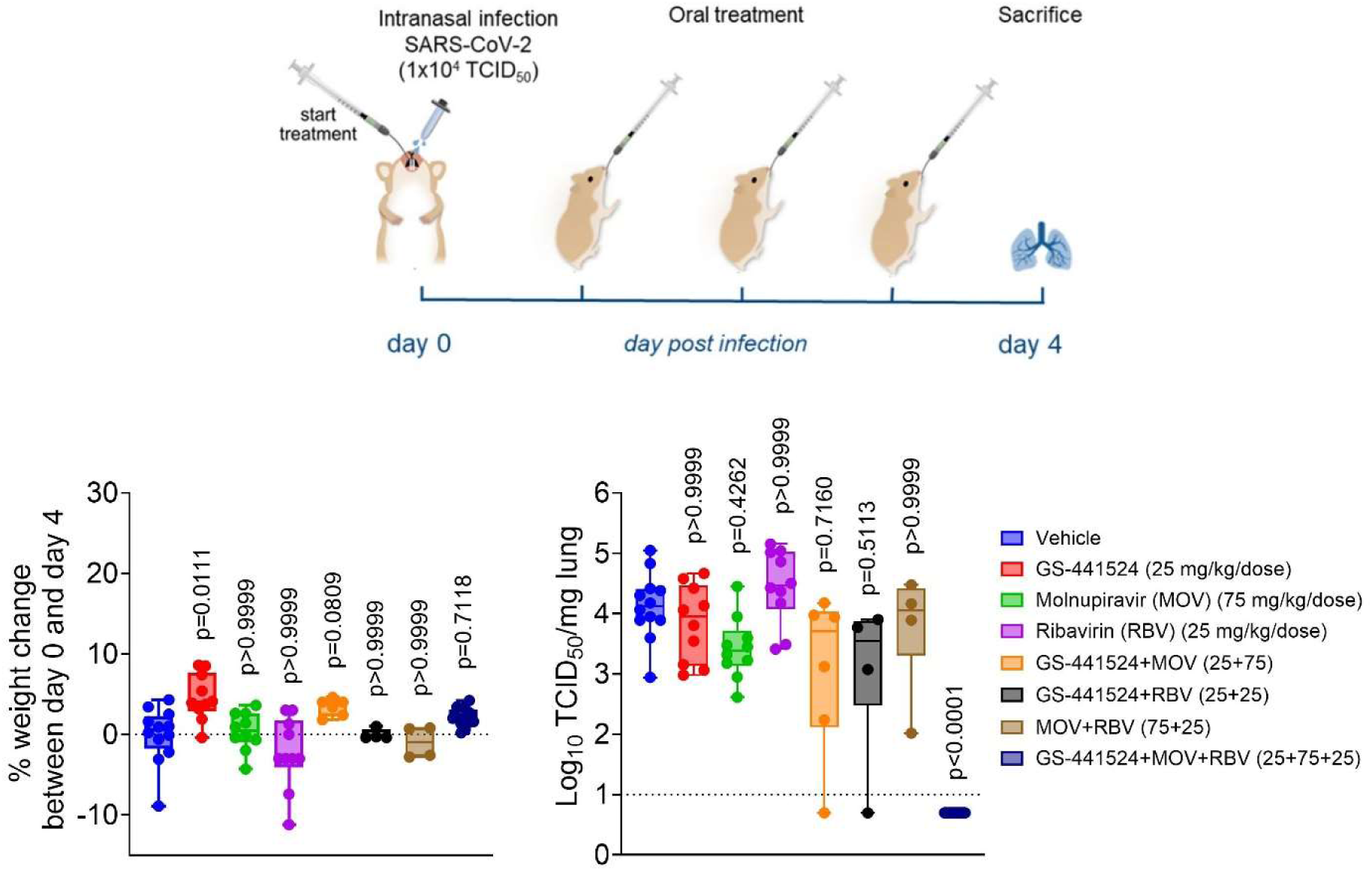
Antiviral effect of combinations of GS-441524, molnupiravir (MOV), and ribavirin (RBV) on SARS-CoV-2 infection in hamsters. (Top) Design of the study. (Bottom) Box plots, with whiskers indicating min to max, representing body weight change (left), and infectious viral titers in the lungs (right). Weight change shown as percentage of change on total weight between day 0 and day 4 p.i.. Infectious virus titers in the lungs at day 4 p.i. as determined as log_10_ TCID_50_ per mg lung tissue. Statistical analysis was performed with the Kruskal-Wallis test with Dunn’s comparison. Data are pool of three independent experiments with n=12 for vehicle and the triple therapy-treated group, n=10 for the monotherapy-treated groups and n=4 or 6 for the dual therapy groups.

## DISCUSSION

Remdesivir and molnupiravir, both prodrugs of nucleoside analogues were initially developed for the treatment of other indications (treatment of Ebola and influenza infections respectively) but were during the SARS-CoV-2 pandemic “repurposed” for the treatment of COVID-19. Besides these nucleoside analogues, potent coronavirus main protease (3CLpro) inhibitors have been developed (nirmatrelvir, ensitrelvir) or are in development. So far, fortunately, the development of resistance of SARS-CoV-2 to the above-mentioned drugs has been limited. However, in such case that during future outbreaks with (a) novel coronavirus(es), such antivirals would be more widely used (in particular during the first phases of a pandemic when vaccines are not yet available), drug resistance may possibly become an issue. Combining drugs with a different resistance profile may then delay or avoid the emergence of such variants. Hence, we believe that it might be prudent to explore the combination of existing drugs against coronaviruses in general and SARS-CoV-2 in particular. As a first step, we aimed to explore the antiviral potency of some combinations and then in a next stage (but not part of this study) explore whether such combinations also prevent the potential development of resistance. To that end, we studied the combination of remdesivir (its parent nucleoside) and molnupiravir but explored in addition whether ribavirin (which is not known to be a coronavirus inhibitor) may further modulate the combined activity of these molecules. The latter was inspired by the fact that ribavirin is, as monotherapy, not active against in the treatment of infections with the hepatitis C virus; yet markedly potentiates the antiviral efficacy of interferon alfa. Furthermore, these three molecules were selected because they each exert activity against families of viruses beyond the coronaviruses (such as para- and orthomyxoviruses); hence the hope that the data from this study may be relevant for the setup of similar studies with yet other viruses.

We first explored the antiviral activity of triple therapy in nasal human airway epithelial cell (HAEC) cultures grown at the air liquid interface infected with HCoV-OC43. Concentrations of each of the drugs were selected such that they had alone only a minimal effect on viral replication. The triple combination of each of these drugs used at suboptimal concentrations, resulted in a pronounced antiviral effect, whereas the double combinations did not. This synergy may be explained by the complementary mechanisms of antiviral action of these molecules. First, ribavirin 5’-monophosphate (RMP - an intracellular metabolite of ribavirin) acts as a competitive inhibitor of inosine 5’-monophosphate dehydrogenase (IMPDH), leading to the depletion of (intracellular) guanosine (deoxy)nucleoside triphosphate (GTP) pools. As GTP is an essential building block for viral RNA synthesis, this consequently inhibits the replication of RNA viruses (24). Decrease in GTP levels limits the concentration of the ATP pools and *vice versa* (25), facilitating the incorporation of other purine nucleoside analogs into nascent RNA such as GS-441524 – an adenosine analogue – which act as a chain terminator. As a result, ribavirin may increase the activity of GS-441524 that ultimately prevents RNA strand elongation and viral replication. Similar synergistic interactions between ribavirin and other purine nucleoside analogues have been reported in Lassa virus and HIV infections (26–28). Second, the active intracellular molnupiravir metabolite, β-D-*N*^4^-hydroxycytidine-triphosphate (NHC-TP), is incorporated into new synthesized viral RNA strands to replace either CTP or UTP, thereby causing C-to-U and G-to-A transition mutations which results in error catastrophe and lethal mutagenesis (7, 29). Therefore, the combined actions of error induction (molnupiravir), chain termination (GS-441524) with depletion of GTP levels (ribavirin) may explain (also the combination of ribavirin and GS-441524) the efficacious suppressing of coronavirus replication.

Inspired by the observations with HCoV-OC43 in nasal HAEC cultures, we explored whether the triple combinations of the three drugs, each used at suboptimal doses, would also result in the SARS-CoV-2 hamster infection model in a pronounced antiviral activity. Whereas each of the drugs alone reduced only minimally (GS-441524 0.2 log_10_ and molnupiravir 0.7 log_10_) or not (ribavirin) infectious viral titers in the lungs; when they were combined, no infectious virus was detectable in the lungs of any of the 12 treated hamsters (> 3.7 log_10_ reduction). The modest antiviral effect of molnupiravir alone at (75 mg/kg/dose) is consistent with data from other studies in hamsters (19, 30, 31). This inhibition is associated with the increased mutation frequency of viral genome (19, 31). The 75 mg/kg molnupiravir dose is equivalent to 10 mg/kg of the Human Equivalent Dose (i.e. 600 mg) when normalised to the human body surface area (32). This is less than the standard dosing regimen of molnupiravir in outpatients with COVID-19 (800 mg, BID) (33). Also, no statistically significant antiviral potency was observed when hamsters were treated with 25 mg/kg/dose GS-441524, which is in agreement with previous report (15). Ribavirin is not known as an inhibitor of coronaviruses neither *in vitro* nor *in vivo* (8, 9, 34–36). A possible explanation is the excision of ribavirin-5’monophosphate after incorporation by the coronavirus exoribonuclease (37). Ribavirin can, however, increase the activity of other antivirals as demonstrated in HCV-, SARS-CoV-, and SARS-CoV-2-infected patients (9, 38, 39). Here we used a dose of 25 mg/kg BID ribavirin which is equivalent to 200 mg BID Human Equivalent Dose (32). This is less than the standard dose of 800 mg/day in the treatment of HCV infections. If our finding could be translated to the human scenario, it is presumable that the triple therapy may results in a potent antiviral effect at clinically readily achievable doses of remdesivir, molnupiravir, and ribavirin with possibly a lower probability of associated adverse events. An important aspect, and still to be explored, is whether this triple combination may also prevent the development of possible drug-resistant variants.

In conclusion, we demonstrate that when suboptimal to even almost inactive doses of the nucleoside analogues GS-441524, molnupiravir, and ribavirin are combined, this results in a pronounced antiviral potency which is most remarkable in a SARS-CoV-2 hamster infection model. Since these and some other small molecule antivirals exert also antiviral activity against viruses belonging to other families, it may in the context of epidemic and pandemic preparedness be important to explore whether the triple combination studied here, or yet other combinations, may hold promise for the treatment of life-threatening viral infections. Also, for those viral infections for which there is today no treatment available, it may be explored whether certain combinations hold promise.

## Author contributions

J.N., D.J., R.A., S.C., T.N.D.D. conceptualized the project; T.N.D.D. and R.A. designed the research; T.N.D.D. and R.A. performed experiments, analyzed data, and made figures; T.N.D.D. wrote the manuscript with input from co-authors; J.N., D.J., S.C., B.B., and R.A. supervised the study; D.J., R.A., J.N., S.C., and B.B. edited the manuscript; J.N., D.J., and S.C. acquired the funding.

## Acknowledgements

We thank Caroline De Keyzer, Lindsey Bervoets, Thibault Francken, Birgit Voeten, Stijn Hendrickx, Niels Cremers for their excellent technical assistance. We also thank Epithelix Sàrl for providing MucilAir cultures in this study. This project has received funding from the Covid-19-Fund KU Leuven/UZ Leuven and the COVID-19 call of FWO (G0G4820N), the European Union’s Horizon 2020 research and innovation program under grant agreements No 101003627 (SCORE project). This work was also supported by the Belgian Federal Government for the VirusBank Platform and the National Institutes of Health (NIH) grant No: R01AI177512-01. T.N.D.D received the fellowship from the European Union’s Horizon 2020 research and innovation programme under Marie Sklodowska-Curie grant agreement No. 812673 (OrganoVIR project).

---

**Figure S1.**
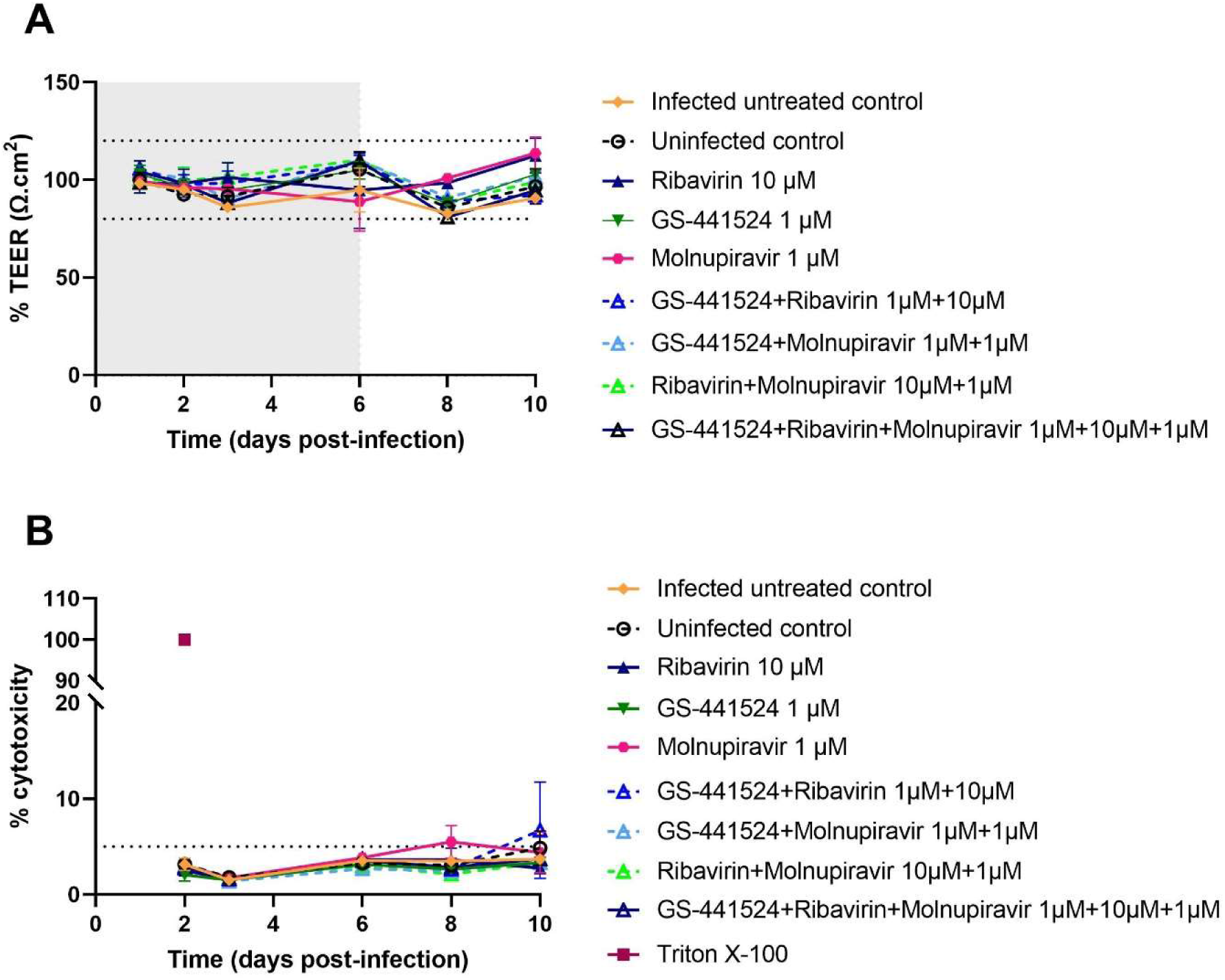
Effect of compound/virus-induced HAE tissue disruption in the presence of dual or triple combinations (data linked to the experiment presented in Figure 2). (A) TEER in response to HCoV-OC43 infection in the presence or absence of compounds at different time points. (B) Time course of HCoV-OC43-induced tissue integrity changes represented by cytotoxicity percentage. The threshold value of 5% cytotoxicity corresponds to a physiological cell turnover in MulcilAir long-term culture (17).

## Reference

1. Halford B. The path to Paxlovid How Pfizer scientists transformed an old drug lead into an oral COVID-19 antiviral. Chemical & Engineering News. 2022:16–8.

2. Li G, Hilgenfeld R, Whitley R, De Clercq E. Therapeutic strategies for COVID-19: progress and lessons learned. Nature Reviews Drug Discovery. 2023.

3. Kabinger F, Stiller C, Schmitzová J, Dienemann C, Kokic G, Hillen HS, et al. Mechanism of molnupiravir-induced SARS-CoV-2 mutagenesis. Nat Struct Mol Biol. 2021;28(9):740–6.

4. Toussi SS, Hammond JL, Gerstenberger BS, Anderson AS. Therapeutics for COVID-19. Nat Microbiol. 2023;8(5):771–86.

5. Mackman RL, Kalla RV, Babusis D, Pitts J, Barrett KT, Chun K, et al. Discovery of GS-5245 (Obeldesivir), an Oral Prodrug of Nucleoside GS-441524 That Exhibits Antiviral Efficacy in SARS-CoV-2-Infected African Green Monkeys. J Med Chem. 2023;66(17):11701–17.

6. Painter GR, Natchus MG, Cohen O, Holman W, Painter WP. Developing a direct acting, orally available antiviral agent in a pandemic: the evolution of molnupiravir as a potential treatment for COVID-19. Curr Opin Virol. 2021;50:17–22.

7. Syed YY. Molnupiravir: First Approval. Drugs. 2022;82(4):455–60.

8. Eslami G, Mousaviasl S, Radmanesh E, Jelvay S, Bitaraf S, Simmons B, et al. The impact of sofosbuvir/daclatasvir or ribavirin in patients with severe COVID-19. J Antimicrob Chemother. 2020;75(11):3366–72.

9. Hung IF, Lung KC, Tso EY, Liu R, Chung TW, Chu MY, et al. Triple combination of interferon beta-1b, lopinavir-ritonavir, and ribavirin in the treatment of patients admitted to hospital with COVID-19: an open-label, randomised, phase 2 trial. Lancet. 2020;395(10238):1695–704.

10. Nyström K, Wanrooij PH, Waldenström J, Adamek L, Brunet S, Said J, et al. Inosine Triphosphate Pyrophosphatase Dephosphorylates Ribavirin Triphosphate and Reduced Enzymatic Activity Potentiates Mutagenesis in Hepatitis C Virus. J Virol. 2018;92(19).

11. Shyr ZA, Cheng YS, Lo DC, Zheng W. Drug combination therapy for emerging viral diseases. Drug Discov Today. 2021;26(10):2367–76.

12. Abdelnabi R, Foo CS, Kaptein SJF, Zhang X, Langendries L, Vangeel L, et al. Molnupiravir (EIDD-2801) inhibits SARS-CoV-2 replication and enhances the efficacy of favipiravir in a Syrian hamster infection model. bioRxiv. 2021:2020.12.10.419242.

13. Chiba S, Kiso M, Nakajima N, Iida S, Maemura T, Kuroda M, et al. Co-administration of Favipiravir and the Remdesivir Metabolite GS-441524 Effectively Reduces SARS-CoV-2 Replication in the Lungs of the Syrian Hamster Model. mBio. 2021;13(1):e0304421.

14. Jonsdottir HR, Siegrist D, Julien T, Padey B, Bouveret M, Terrier O, et al. Molnupiravir combined with different repurposed drugs further inhibits SARS-CoV-2 infection in human nasal epithelium in vitro. Biomed Pharmacother. 2022;150:113058.

15. Abdelnabi R, Maes P, de Jonghe S, Weynand B, Neyts J. Combination of the parent analogue of remdesivir (GS-441524) and molnupiravir results in a markedly potent antiviral effect in SARS-CoV-2 infected Syrian hamsters. Front Pharmacol. 2022;13:1072202.

16. Tapparel C, Sobo K, Constant S, Huang S, Van Belle S, Kaiser L. Growth and characterization of different human rhinovirus C types in three-dimensional human airway epithelia reconstituted in vitro. Virology. 2013;446(1-2):1–8.

17. Boda B, Benaoudia S, Huang S, Bonfante R, Wiszniewski L, Tseligka ED, et al. Antiviral drug screening by assessing epithelial functions and innate immune responses in human 3D airway epithelium model. Antiviral Res. 2018;156:72–9.

18. Abdelnabi R, Foo CS, Jochmans D, Vangeel L, De Jonghe S, Augustijns P, et al. The oral protease inhibitor (PF-07321332) protects Syrian hamsters against infection with SARS-CoV-2 variants of concern. Nat Commun. 2022;13(1):719.

19. Abdelnabi R, Foo CS, Kaptein SJF, Zhang X, Do TND, Langendries L, et al. The combined treatment of Molnupiravir and Favipiravir results in a potentiation of antiviral efficacy in a SARS-CoV-2 hamster infection model. EBioMedicine. 2021;72:103595.

20. Reed LJ, Muench H. A SIMPLE METHOD OF ESTIMATING FIFTY PER CENT ENDPOINTS. American Journal of Epidemiology. 1938;27(3):493–7.

21. Kaptein SJF, Jacobs S, Langendries L, Seldeslachts L, Ter Horst S, Liesenborghs L, et al. Favipiravir at high doses has potent antiviral activity in SARS-CoV-2-infected hamsters, whereas hydroxychloroquine lacks activity. Proc Natl Acad Sci U S A. 2020;117(43):26955–65.

22. Poch G, Dittrich P, Holzmann S. Evaluation of combined effects in dose-response studies by statistical comparison with additive and independent interactions. J Pharmacol Methods. 1990;24(4):311–25.

23. Fuchs EJ, Kiser JJ, Hendrix CW, Sulkowski M, Radebaugh C, Bushman L, et al. Plasma and intracellular ribavirin concentrations are not significantly altered by abacavir in hepatitis C virus-infected patients. J Antimicrob Chemother. 2016;71(6):1597–600.

24. Paeshuyse J, Dallmeier K, Neyts J. Ribavirin for the treatment of chronic hepatitis C virus infection: a review of the proposed mechanisms of action. Curr Opin Virol. 2011;1(6):590–8.

25. Pelley JW. 14 -Purine, Pyrimidine, and Single-Carbon Metabolism. In: Pelley JW, editor. Elsevier’s Integrated Review Biochemistry (Second Edition). Philadelphia: W.B. Saunders; 2012. p. 119–24.

26. Margot NA, Miller MD. In vitro combination studies of tenofovir and other nucleoside analogues with ribavirin against HIV-1. Antivir Ther. 2005;10(2):343–8.

27. Oestereich L, Rieger T, Ludtke A, Ruibal P, Wurr S, Pallasch E, et al. Efficacy of Favipiravir Alone and in Combination With Ribavirin in a Lethal, Immunocompetent Mouse Model of Lassa Fever. J Infect Dis. 2016;213(6):934–8.

28. Hartman NR, Ahluwalia GS, Cooney DA, Mitsuya H, Kageyama S, Fridland A, et al. Inhibitors of IMP dehydrogenase stimulate the phosphorylation of the anti-human immunodeficiency virus nucleosides 2’,3’-dideoxyadenosine and 2’,3’-dideoxyinosine. Mol Pharmacol. 1991;40(1):118–24.

29. Urakova N, Kuznetsova V, Crossman DK, Sokratian A, Guthrie DB, Kolykhalov AA, et al. β-D-N4-Hydroxycytidine Is a Potent Anti-alphavirus Compound That Induces a High Level of Mutations in the Viral Genome. Journal of Virology. 2018;92(3):e01965–17.

30. Jeong JH, Chokkakula S, Min SC, Kim BK, Choi WS, Oh S, et al. Combination therapy with nirmatrelvir and molnupiravir improves the survival of SARS-CoV-2 infected mice. Antiviral Res. 2022;208:105430.

31. Sheahan TP, Sims AC, Zhou S, Graham RL, Pruijssers AJ, Agostini ML, et al. An orally bioavailable broad-spectrum antiviral inhibits SARS-CoV-2 in human airway epithelial cell cultures and multiple coronaviruses in mice. Sci Transl Med. 2020;12(541).

32. Reagan-Shaw S, Nihal M, Ahmad N. Dose translation from animal to human studies revisited. Faseb j. 2008;22(3):659–61.

33. Jayk Bernal A, Gomes da Silva MM, Musungaie DB, Kovalchuk E, Gonzalez A, Delos Reyes V, et al. Molnupiravir for Oral Treatment of Covid-19 in Nonhospitalized Patients. N Engl J Med. 2022;386(6):509–20.

34. Al-Tawfiq JA, Momattin H, Dib J, Memish ZA. Ribavirin and interferon therapy in patients infected with the Middle East respiratory syndrome coronavirus: an observational study. Int J Infect Dis. 2014;20:42–6.

35. Day CW, Baric R, Cai SX, Frieman M, Kumaki Y, Morrey JD, et al. A new mouse-adapted strain of SARS-CoV as a lethal model for evaluating antiviral agents in vitro and in vivo. Virology. 2009;395(2):210–22.

36. Zandi K, Amblard F, Musall K, Downs-Bowen J, Kleinbard R, Oo A, et al. Repurposing Nucleoside Analogs for Human Coronaviruses. Antimicrob Agents Chemother. 2020;65(1).

37. Ferron F, Subissi L, Silveira De Morais AT, Le NTT, Sevajol M, Gluais L, et al. Structural and molecular basis of mismatch correction and ribavirin excision from coronavirus RNA. Proc Natl Acad Sci U S A. 2018;115(2):E162–E71.

38. Chu CM, Cheng VC, Hung IF, Wong MM, Chan KH, Chan KS, et al. Role of lopinavir/ritonavir in the treatment of SARS: initial virological and clinical findings. Thorax. 2004;59(3):252–6.

39. Jen J, Laughlin M, Chung C, Heft S, Affrime MB, Gupta SK, et al. Ribavirin dosing in chronic hepatitis C: application of population pharmacokinetic-pharmacodynamic models. Clin Pharmacol Ther. 2002;72(4):349–61.

40. Tao K, Tzou PL, Nouhin J, Bonilla H, Jagannathan P, Shafer RW. SARS-CoV-2 Antiviral Therapy. Clin Microbiol Rev. 2021;34(4):e0010921.

41. Vehreschild M, Atanasov P, Yurko K, Oancea C, Popov G, Smesnoi V, et al. Safety and Efficacy of Vidofludimus Calcium in Patients Hospitalized with COVID-19: A Double-Blind, Randomized, Placebo-Controlled, Phase 2 Trial. Infect Dis Ther. 2022;11(6):2159–76.

42. Schultz DC, Johnson RM, Ayyanathan K, Miller J, Whig K, Kamalia B, et al. Pyrimidine inhibitors synergize with nucleoside analogues to block SARS-CoV-2. Nature. 2022;604(7904):134–40.

